# HAIRpred: Prediction of human antibody interacting residues in an antigen from its primary structure

**DOI:** 10.1101/2024.12.17.628825

**Authors:** Ruchir Sahni, Nishant Kumar, Gajendra P. S. Raghava

**Author notes:** **Mailing Address of Authors** Ruchir Sahni (RS) Nishant Kumar (NK) Gajendra P. S. Raghava (GPSR). **Corresponding Author** Prof. Gajendra P. S. Raghava, Head and Professor, Department of Computational Biology, Indraprastha Institute of Information Technology, Delhi, Okhla Industrial Estate, Phase III (Near Govind Puri Metro Station), New Delhi, India – 110020 Office: A-302 (R&D Block), Website: http://webs.iiitd.edu.in/raghava/.

## Abstract

1.

In the past, several methods have been developed for predicting conformational B-cell epitopes in antigens that are not specific to any host. Our primary analysis of antibody-antigen complexes indicated a need to develop host-specific B-cell epitopes. In this study, we present a novel approach to predict conformational B-cell epitopes specific to human hosts by focusing on human antibody-interacting residues in antigens. We trained, test and evaluate our models on 277 complexes of human antibody-antigen complexes. Initially, we employed machine learning models based on the binary sequence profile of antigens, achieving a maximum area under the receiver operating characteristic curve (AUC) of 0.61. Performance of model improved significantly AUC from 0.61 to 0.67, when evolutionary profiles are used instead of binary profiles. Models developed using embeddings from fine-tuned large language models reached an AUC of 0.61. Additionally, models utilizing predicted surface relative solvent accessibility achieved an AUC of 0.67. Our ensemble model, which combined relative surface accessibility with evolutionary profiles, achieved highest precision with an AUCROC of 0.72. All models in this study were trained using five-fold cross-validation on a training dataset and evaluated on an independent dataset not used for training or testing. Our method outperforms existing approaches on the independent dataset. Furthermore, we used SHAP eXplainable AI (XAI) method to interpret the importance of individual features contributing to the predictions made by our models. To support the scientific community, we have developed a standalone software, and web server, HAIRpred, for predicting human antibody-interacting residues in proteins. https://webs.iiitd.edu.in/raghava/hairpred/.

**Author’s Biography:**

1. Ruchir Sahni is currently studying as an integrated BS-MS student at Indian Institute of Science Education and Research (IISER) Pune, India. He is currently working as an Intern on Project position at Department of Computational Biology, Indraprastha Institute of Information Technology (IIIT), New Delhi, India.
2. Nishant Kumar is currently working as Ph.D. in Computational biology from Department of Computational Biology, Indraprastha Institute of Information Technology (IIIT), New Delhi, India.
3. Gajendra P. S. Raghava is currently working as Professor and Head of Department of Computational Biology, Indraprastha Institute of Information Technology (IIIT), New Delhi, India.

## 2. Introduction

Antibodies are specialized glycoproteins produced by the immune system, playing a vital role in identifying and neutralizing pathogens, as well as in establishing immunological memory. Their essential functions make antibodies crucial for maintaining human health and supporting immune responses. Given their wide-ranging applications in diagnostics, vaccines, immunotherapy, and therapeutics, the study of antibody activity and interactions is a fundamental aspect of immunological research. The interaction between an antibody and its corresponding antigen predominantly occurs through specific surface residues on the antigen, termed antibody-interacting residues. These residues are critical in defining conformational B-cell epitopes, as they represent the direct contact points between antibodies and their targets. Accurately identifying these residues is therefore essential for predicting conformational B-cell epitopes, which play a significant role in immune recognition. Conventional experimental techniques for identifying antibody-interacting regions, such as X-ray crystallography and alanine scanning mutagenesis are labour-intensive, costly, and often unsuitable for large-scale analysis [1].

Recent progress in machine learning and artificial intelligence has led to the development of various computational models aimed at predicting antibody-interacting residues or conformational B-cell epitopes. Broadly, existing methods for predicting conformational B-cell epitopes can categorized in two groups namely structure-based and sequence-based methods. Following are examples of structure-based methods commonly used for predicting conformational B-cell epitopes in an antigen from its tertiary structure CEP, DiscoTope, PEPITO, Epitopia, SEPPA, SEMA, and Epitope3D [2–9]. One of the limitation of structure-based methods is that they need tertiary structure of antigen. In order to address this challenge number of sequence-based methods have been developed for predicting conformational B-cell epitopes in an antigen from its amino acid sequence. Following are commonly used sequence-based methods BepiPred-3.0, SEMA-2.0, CBTope, and CLBTope [10–13]. However, a notable limitation of these models is their inability to accommodate host-specific differences in antibody-antigen interactions. Antibodies are finely adapted to their respective host organisms, meaning that the epitopes recognized by human antibodies can differ significantly from those recognized by antibodies from other species, such as mice. Additionally, these models frequently overlook variations in immune responses, antibody structures, and post-translational modifications across different species [14,15]. Such limitations raise concerns about the generalizability of these models across datasets obtained from diverse experimental settings. Recent research by Cia et al. (2023) has highlighted the shortcomings of existing methods by demonstrating their limited performance when tested against experimentally derived complexes from the Protein Data Bank (PDB) [16]. As a result, focusing on human antibody-interacting residues to predict epitopes that are specific to human antibodies is a crucial step toward enhancing the accuracy and relevance of immune response predictions.

In this study, we present HAIRpred, a novel computational tool designed to predict human antibody-interacting regions in antigens with improved accuracy and interpretability. Our models were developed and validated using experimentally derived human antibody-antigen complexes, ensuring a focus on host specificity. We utilized a carefully curated set of sequence-derived features that have shown predictive efficacy in prior studies while ensuring biological relevance. This feature set integrates structural, sequential, and evolutionary information about the antigen that can be extracted from its sequence. The performance of HAIRpred was rigorously evaluated against existing methods using a diverse dataset of experimentally verified antibody-antigen complexes. Additionally, SHAP method was used to interpret the models which provided an insight into relative contribution of each feature to the prediction outcomes.

## 3. Materials and Methods

### 3.1. Creation of Datasets

We obtained antibody-antigen complexes for human host from SAdDab database (Dunbar et al., 2013) [17]. Antigens or proteins having resolution less than 3.0 Å and an R-factor less than 0.30 were selected for further processing. All antigens having less than 50 residues were removed from dataset. This resulted in a set of 1620 human antibody-antigen complexes. In a similar way, we obtained 95 mouse antibody-antigen complexes from the SAbDaB database. In order to remove redundancy from antigens recognized by human antibodies, we clustered these antigens using CD-HIT with 70% cut-off [18]. Our dataset contain 277 antigens, where no two antigen have more than 70% similarity with each other. This dataset were further divided in training and independent dataset (80:20 ratio), where training dataset contain 221 and independent dataset contain 56 antigens. In order to assign antibody interacting residues in these antigens, we examined changed in relative solvent accessible surface area (RSA) of these residues with and without antibody binding, similar to the algorithm performed by Cia et al. (2023). A residue is assigned antibody interacting residues if it undergo a change in RSA of at least 5% upon binding with an antibody (RSA_unbound_ − RSA_bound_ ≥ 5%) [16].

### 3.2. Creation of overlapping patterns

Numerous studies have shown that the residue’s function is influenced by its local environment [13,19,20]. In this method, we predict antibody interacting and non-interacting residues in an antigen. To capture the local environment of each residue, we created overlapping patterns (or windows) for each residue in the antigen with window sizes ranging from 13 to 21. To generate a pattern for the terminal residues, we added a dummy variable ‘X’ at both ends of antigen. Each pattern’s corresponding label (antibody interacting/not interacting) is the label of the central residue. This approach has been commonly used in multiple studies [21,22]. Patterns were created for both the training dataset and the independent test dataset. The patterns generated for the training dataset were imbalanced i.e. the not-interacting patterns are much larger in number. To avoid any bias due to the imbalance, the balanced training dataset was made which consisted of all antibody interacting patterns and an equal number of randomly selected not-interacting patterns.

### 3.3. Compositional Analysis

Amino acid composition (AAC) is the frequency of each of the 20 amino acids and one dummy residue within a peptide or protein sequence. It is represented as a feature vector with 21 elements, where each component corresponds to the fraction of a particular amino acid residue within the sequence [23]. It can be calculated by equation 1.

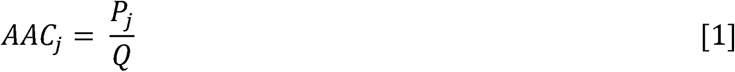

where AACj is amino acid composition of residue type j; Pj and Q number of residues of type j and length of sequence, respectively

### 3.4. Sequence Logo

In order to show sequence conservation in antibody interacting and non-interacting patterns, we generated the sequence logo using a web-based application called “Two Sample Logo”. This tool calculates the significance of each residue at each position and compares distributions between positive and negative samples. Residues are grouped based on enrichment or depletion in the positive sample, and the graphical output can display statistically significant residues with symbol sizes proportional to the difference between the groups. The p value is calculated using t-test [24]. To generate the Two Sample Logo, we inputted all the antibody interacting patterns in the positive sample. To get a better understanding, we only considered those antibody not-interacting patterns in which no residue was antibody interacting. From these patterns, we then randomly selected the negative sample to have the total number of patterns to be equal in both positive and negative sample.

### 3.5. Curation of Feature Set

Most of the existing machine learning techniques need input in form of a numerical vector or matrices. Thus, we need to generate features which would represent the interacting and non-interacting patterns of antigen in the form of a numerical vector. In this study, we curated a wide range of features that capture structural, sequential and evolutionary information of antigen from its sequence. The curated set of features are Binary Profiles, Protein Language Model (PLM) Embeddings, Position-Specific Scoring Matrices (PSSM), predicted Relative Surface Accessibility (RSA) and Secondary Structure (SS).

Binary Profiles were generated by assigning binary values to each amino acid. This results in a 20-length vector for each residue. For example, Arginine (A) was represented as [1,0,0,0,0,0,0,0,0,0,0,0,0,0,0,0,0,0,0,0] and the dummy variable X is represented as a 20 length zero vector. Binary Profiles were created for each pattern, resulting in the dimension of the feature vector as Nx20, where N is the pattern length [13].

Position-Specific Scoring Matrices (PSSM) are matrices containing information about the probability of amino acids occurring in each position. It is derived from a multiple sequence alignment and it represents the evolutionary history of the protein. Here, we had generated PSSM profiles of a sequence using Position-Specific Iterative Basic Local Alignment Search Tool (PSI-BLAST) [25] and searching against the Swiss Prot database [26]. The search was performed with an E-value threshold of 0.1 and a word size of 3 for protein sequences in 3 iterations. The alignment was conducted using BLOSUM62 as the scoring matrix with gap penalties of values of 11 (gap opening) and 1 (gap extension). We generated PSSM profiles for each antigen in the complex and then divided them into patterns. The dummy variable X is assigned a 20 length zero vector while creating the feature vector. The resulting feature vector is a matrix of dimensions Nx20, where N is the length of the pattern.

Protein Language Models (PLM) are transformer based models which generate embeddings that capture contextual relationships between amino acids, reflecting evolutionary and functional information. Here we use ESM2 (esm2_t6_8M_UR50D) [27] and Encoder only ProtTrans (ProtT5-XL-UniRef50) [28] to generate embeddings for the patterns. The patterns are inputted into the models and the last layer embeddings are extracted as a feature vector.

Relative Surface Accessibility (RSA) gives a measure of the extent of exposure of a residue in the 3D structure. To predict RSA from sequence, the 3D structure was predicted using ESMFold [27] and passed through the DSSP algorithm [29] to get RSA values for each residue. These predicted RSA values were then divided into patterns corresponding to their respective residues. The dummy variable X is assigned the RSA of value 0. The resulting feature vector is a vector of length N, where N is the length of the pattern.

Secondary Structure (SS) provides information about the local structure of the protein backbone. Secondary Structure was predicted by two ways: 1) from ProtT5-XL-U50 embeddings (8-class SS) [28] 2) from DSSP algorithm applied on 3D structure predicted by ESMFold (3-class SS) [27]. The predicted secondary structures were then divided into patterns corresponding to their respective residues. The dummy variable X is assigned the secondary structure of a coil. The secondary structure is then encoded using a label encoder, resulting in a vector of length N, where N is the length of the pattern.

### 3.6. Machine Learning and Neural Network Models

The machine learning algorithms were implemented using scikit-learn [30] to develop good predictive models. We focused on the Random Forest algorithm and the XGBoost algorithm due to their strong classification performance and ability to handle variable datasets. The hyperparameters of the models were optimised using GridSearchCV from the scikit-learn package. In addition to traditional machine learning algorithms, we implemented neural network models using PyTorch [31]. Neural networks are highly effective for capturing complex patterns in data due to their multi-layered architecture. The neural networks were optimised using backpropagation and the Adam optimizer. Ensemble Models were also created to combine information from different features. We have used a simple method for creating ensemble models : averaging the probabilities generated by the individual models. The averaged probabilities are then used to decide the predicted label.

### 3.7. Optimization and Evaluation of Models

To optimize our classification models, we used the five-fold cross-validation technique. In this approach, the dataset is randomly divided into five equal-sized "folds." The model is trained on four of these folds and evaluated on the remaining fold, with this process repeated five times. The results from each fold are then averaged to provide an estimate of the model’s performance. This method helps optimize the model’s hyperparameters to maximize AUC. Although different folds were used for training and testing, some degree of overoptimization cannot be fully ruled out with five-fold cross-validation. Therefore, we evaluated our final models on an independent dataset that was not used for hyperparameter optimization.

#### 3.7.1. Evaluation Parameters

In this study, the models are performing binary classification on each window to label each residue as antibody interacting/not-interacting. The performance of the models are evaluated based on the standard evaluation parameters. The evaluation parameters can be divided into threshold-dependent and threshold independent parameters. The threshold independent parameter used in this study is the Area Under the Receiver Operating Characteristic (AUROC) and the threshold dependent parameters used are: Sensitivity, Specificity, Accuracy and Matthew’s correlation coefficient (MCC). These parameters are implemented using the scikit-learn package.

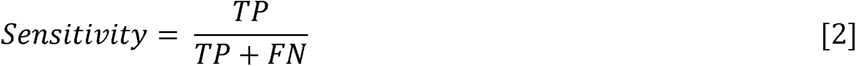

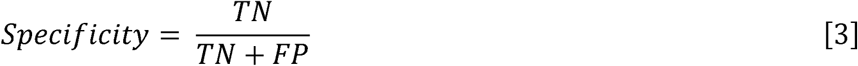

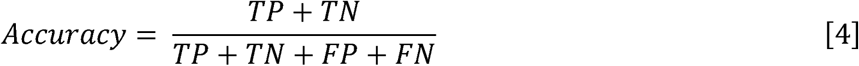

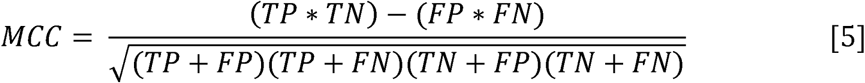

Here TN, TP, FN, and FP stands for true negative, true positive, false negative, and false positive respectively.

### 3.8. Explainable Machine Learning using SHAP

Understanding the decision making process of a given machine learning model is crucial for model reliability, especially in complex biological applications. In this study, we used SHapley Additive exPlanations (SHAP), a post-hoc interpretability method based on Shapley values and cooperative game theory, to analyse machine learning models [32]. We used the Shapley values to ascertain the importance of each feature and the extent of each feature’s contribution to the model’s performance. The Python SHAP package was used for SHAP TreeExplainer to interpret the results of the individual models of the HAIRpred’s Random Forest ensemble.

### 3.9. Web Server

To aid the scientific community, we designed a web server called “HAIRpred” to predict human antibody interacting residues in the antigen (https://webs.iiitd.edu.in/raghava/hairpred/). The user-friendly front end was developed using HTML, CSS, and PHP scripts. The webserver includes the “Predict” module, and the “Design” module.

## 4. Results

We have divided the result section into five categories: (i) Data analysis, (ii) Machine Learning methods, (iii) Ensemble methods, (iv) Benchmarking, and (v) Web Server. The complete workflow of the study is illustrated in Figure 1, and the details of the following subsections can be found below.

**Figure 1:**
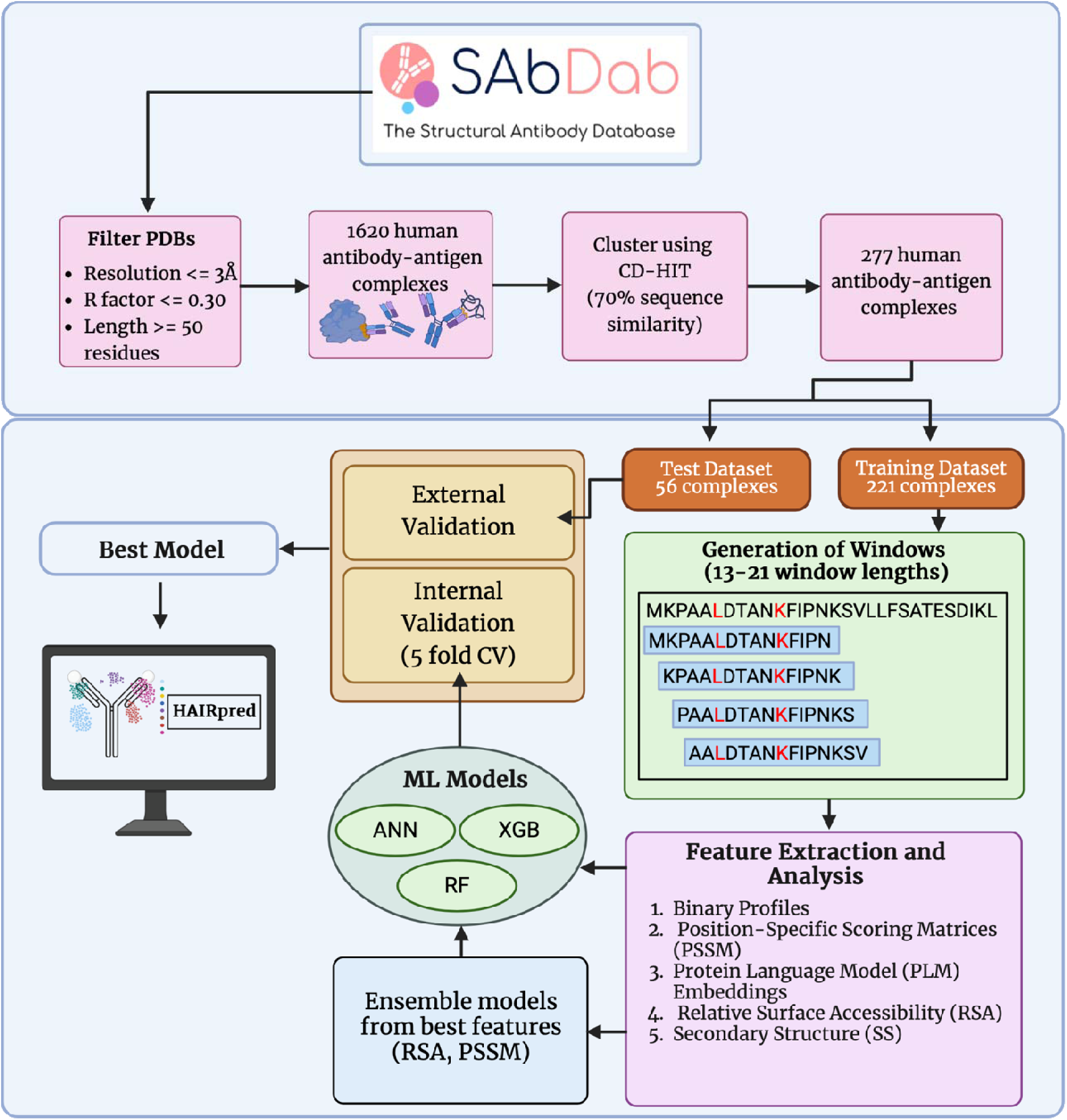
Workflow of the Study

### 4.1. Data Analysis

In the data analysis, we have performed the compositional analysis and two sample logo for antibody interacting and non-interacting residues.

#### 4.1.1. Amino Acid Composition Comparison between Human and Mouse

We compared the Amino Acid Composition (AAC) for both Human and Mouse interacting residues (Figure 3) and general proteome. It was observed that the AAC is significantly different between the two species. For example amino acids like Cysteine (C), Glutamine (Q), Arginine (R), Tryptophan (W), and Tyrosine (Y) are preferred in antibody-antigen interaction in case of human but not in case of mouse. Similarly, residues like Glycine (G), Lysine (K), Asparagine (N) and Serine (S) are preferred in mouse antibody interaction but not in human antibody interaction. This result underscores the importance of host specificity and emphasises on the need for host-specific predictors.

**Figure 3:**
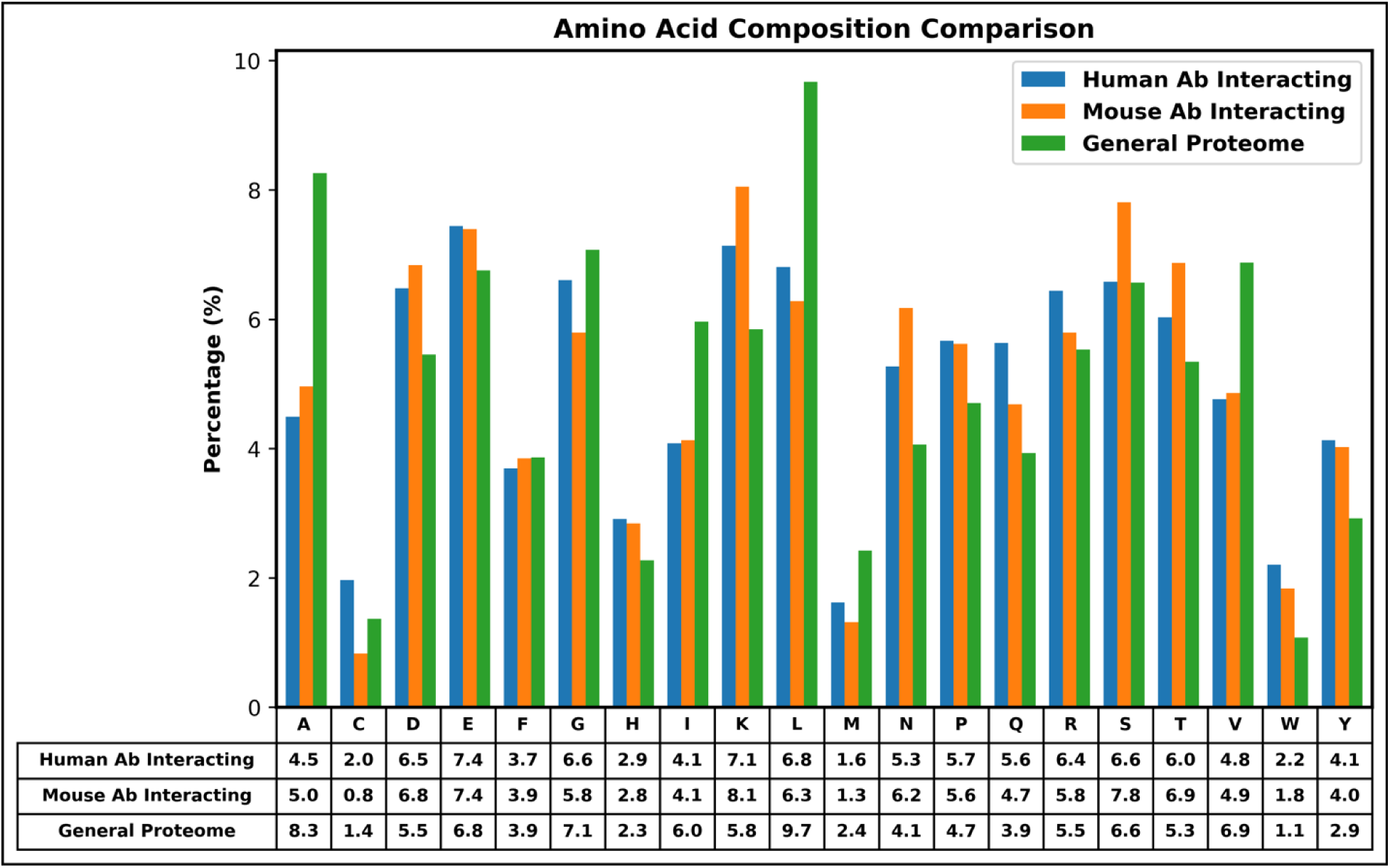
Comparison of amino acid composition of antigens that interact with human and mouse antibodies. It also include amino acid composition of general proteome to provide comparison with base.

#### 4.1.2. Analysis of Human Interacting Residues

We compared the amino acid composition of antibody interacting residues and not-interacting residues (Figure 4) to understand preference of residues. The p-values are generated using the chi-square test. The composition of residues Aspartic Acid (D), Glutamic Acid (E), Histidine (H), Lysine (K), Proline (P), Glutamine (Q), Arginine (R), Tryptophan (W), and Tyrosine (Y) is significantly higher in antibody interacting regions. Similarly, composition of Alanine (A), Cysteine (C), Glycine (G), Isoleucine (I), Leucine (L) and Valine (V) is significantly lower in antibody interacting regions. Antibody interacting regions are characterised by a mix of charged, polar, and aromatic residues whereas not-interacting regions are dominated by hydrophobic and mildly polar residues.

**Figure 4:**
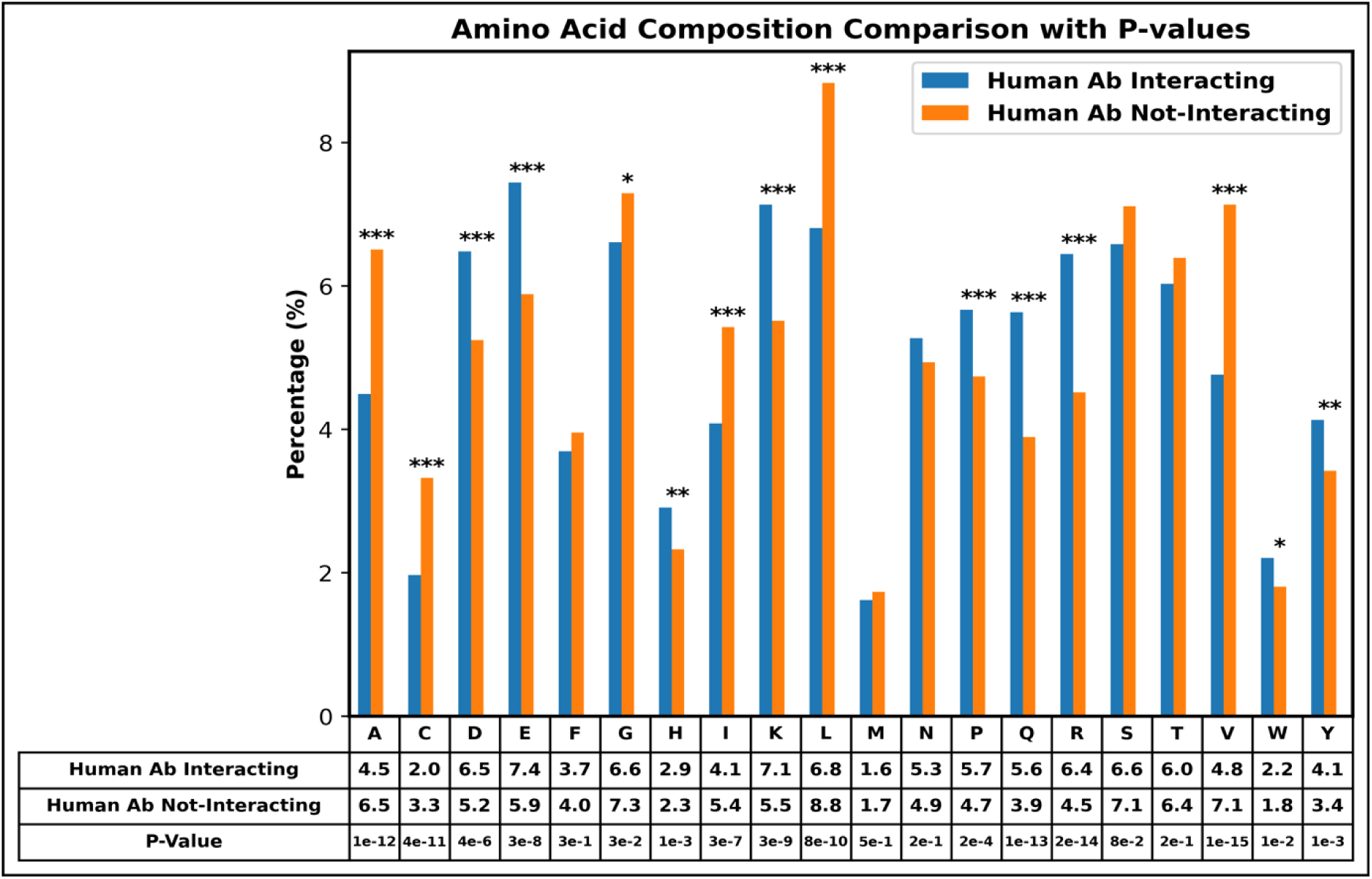
Percentage Composition of Antibody Interacting Residues and Antibody Not-Interacting residues in Antigen

#### 4.1.3. Two Sample Logo

In this study, we have built the two sample logo to understand the preference of a residue at a specific position in human antibody interacting patterns. The two sample logo is displayed in Figure 5. The central residue results are similar to the AAC performed in Figure 4. Alanine (A) is highly abundant in antibody not-interacting residues while in antibody interacting residues, Glutamine (Q) and Cysteine (C) are in abundance.

**Figure 5:**
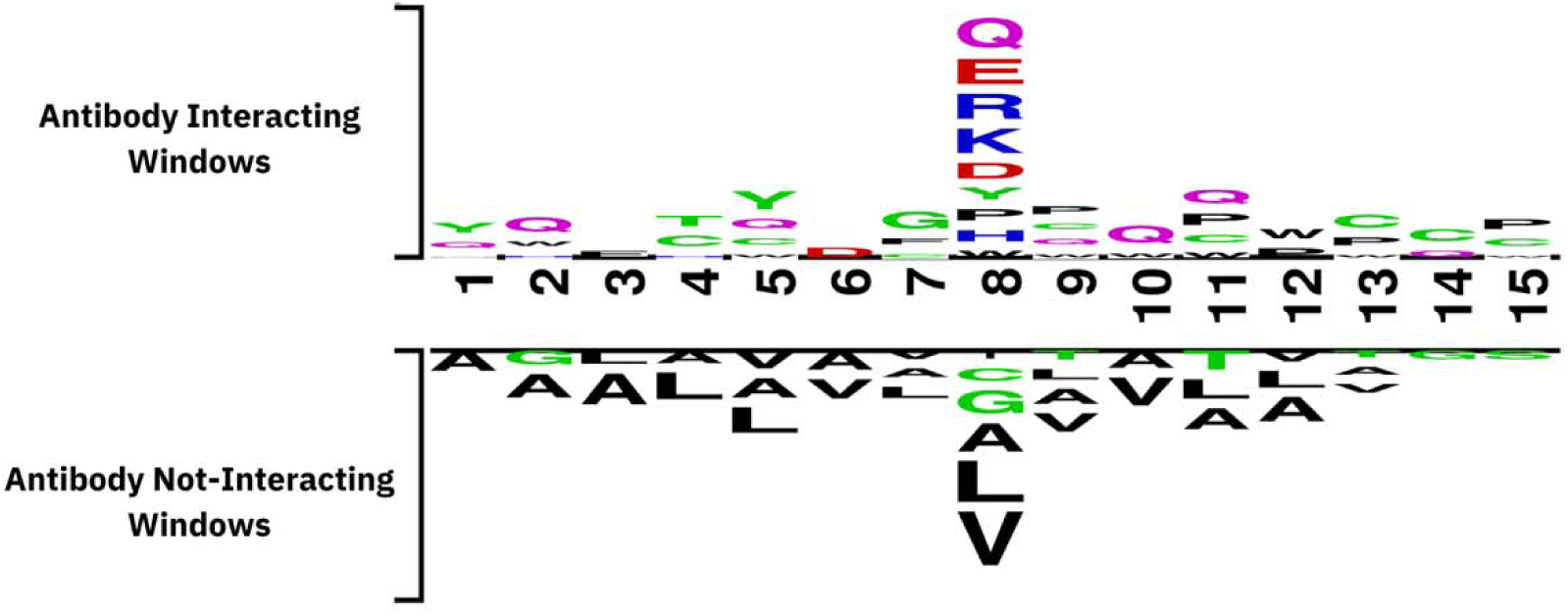
Two Sample Logo constructed from the interacting and non-interacting patterns in the human dataset

### 4.2. Machine Learning Methods

#### 4.2.1. Models developed using sequence and evolutionary features

As most of the machine learning techniques need features in fixed length numerical vectors to develop prediction models. Thus, we generate numerical features corresponding to interacting and non-interacting patterns. It has been shown in number of studies in the past that pattern/window length of seventeen is most effective for predicting of interacting residues [33–35]. In this study, we derived Binary and PSSM Profile for each pattern to extract sequence and evolutionary information of a pattern. These features were used to train the machine learning algorithms and the predictive performance for each model was then evaluated against the independent test dataset. As shown in Table 1, our random forest (RF) achieved maximum performance AUC of 0.61 with MCC 0.11. The performance our models improved significantly, when we used evolutionary information of patterns in form of PSSM profile. As shown in Table 1, our RF-based model obtain highest AUC 0.67 with MCC 0.17. These results agree with previous studies where PSSM based models perform better than sequence based models [19,20].

**Table 1:**
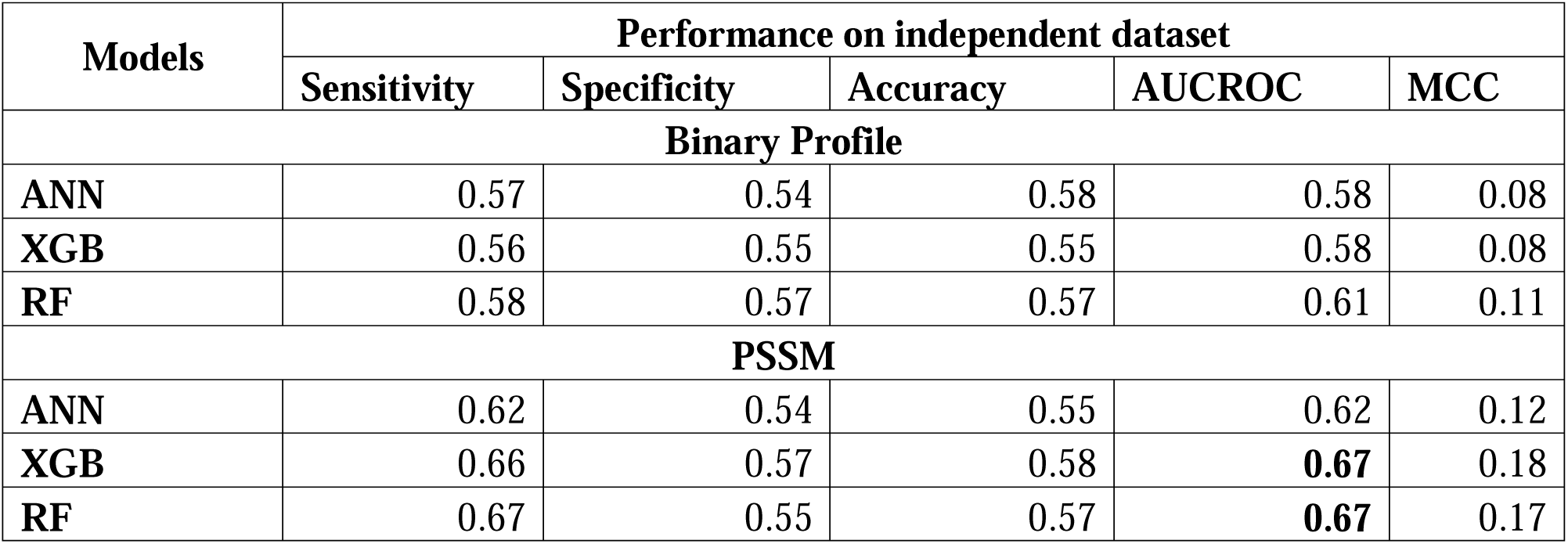
The performance of machine learning based models on independent dataset, developed using binary and PSSM profiles at window length seventeen.

#### 4.2.2. Models developed using LLM embeddings and structural features

In addition to sequence and evolutionary information, we also used embeddings generated by protein-language models (PLMs). Protein language models are based on natural language processing where they generate embeddings from large-scale protein data [36]. In this study, we used PLM embeddings as a feature vector as well as used PLMs to predict structural information including Relative Solvent Accessibility (RSA) and Secondary Structure (SS) which were used as feature vectors. Similar to sequence based features, Each feature vector was generated from the training dataset on the patterns of length seventeen and used to train three models: Random Forest, XGBoost, and Artificial Neural Network. The predictive performance of each feature was then evaluated against the independent test dataset. It is observed that the RSA feature gives the best performance with the AUC score reaching 0.67. The results are displayed in Table 2. Additional data is available in the supplementary table S1.

**Table 2:**
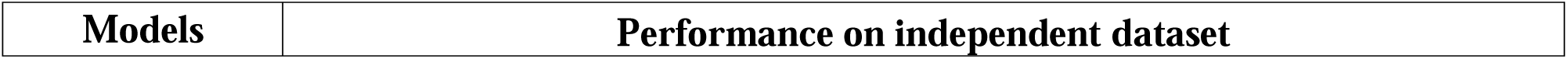
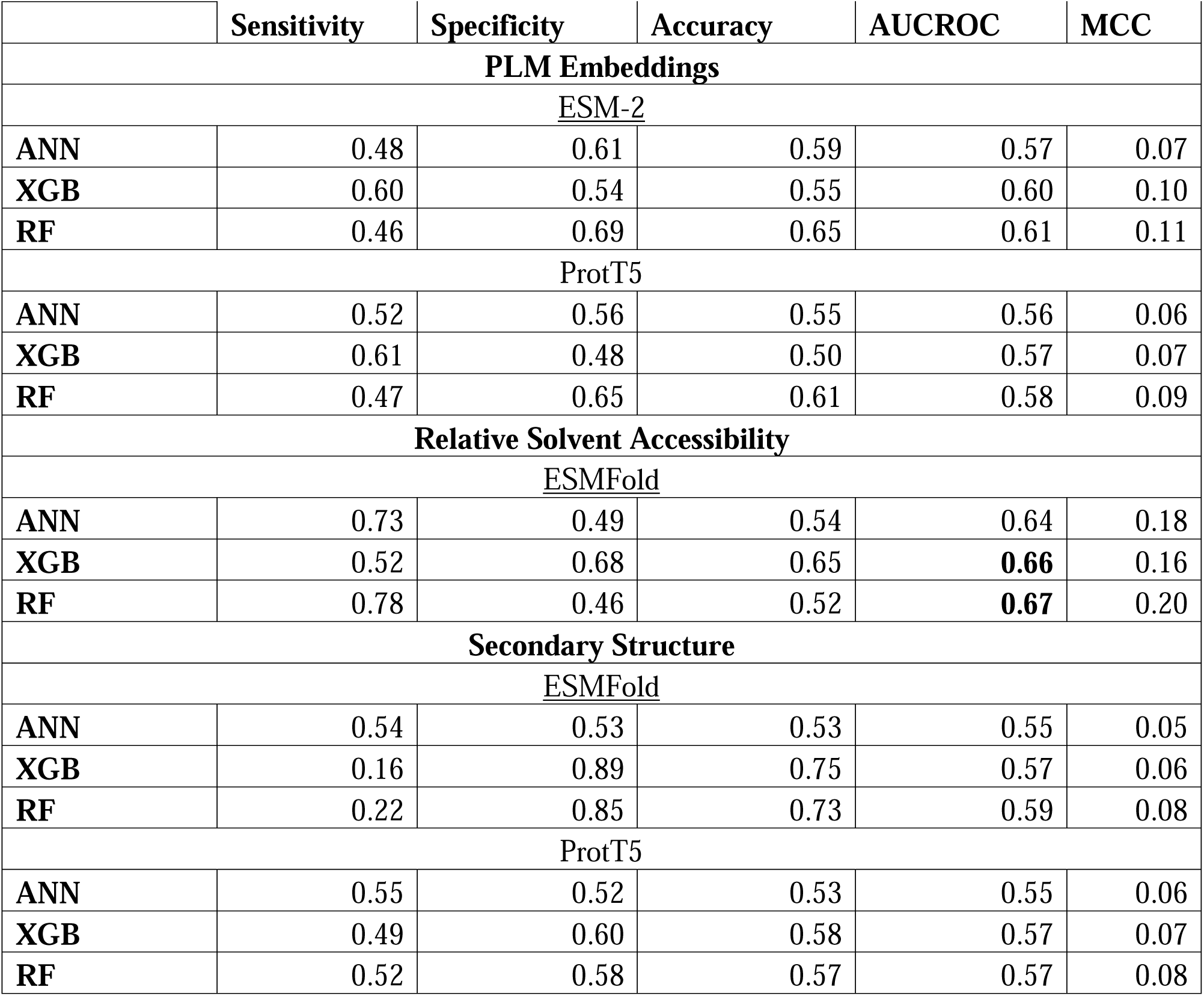
The performance of machine learning models developed using PLM generated embeddings, predicted secondary structure and relative surface accessibility.

### 4.3. Ensemble Models

In our analysis, we observed that models developed using PSSM profile and relative surface accessibility perform better than other models. In order to utilize strength of these feature, we developed an ensemble model that combine both type of features. As shown in Table 3, we achieved highest AUC 0.71 with MCC 0.22 at window length seventeen. So far we used window/pattern length 17, as this most of previous studies used 17. In order to understand effect of window length on the performance of our models. We developed models using window length 13 to 21 and compute the performance of our ensemble models. It was observed that our RF-based model perform best on window length 15 with AUC 0.72 and MCC 0.23. The performance is marginally better for window length 15 then window length 17. Additional data is available in the supplementary table S2.

**Table 3:** The performance of the ensemble model using different window lengths on an independent dataset.

| Models | Metrics evaluated on the Independent Test Dataset |  |  |  |  |
| --- | --- | --- | --- | --- | --- |
|  | Sensitivity | Specificity | Accuracy | AUCROC | MCC |
| **Window Length – 13** |  |  |  |  |  |
| **ANN** | 0.66 | 0.57 | 0.59 | 0.67 | 0.18 |
| **XGB** | 0.72 | 0.54 | 0.57 | 0.70 | 0.20 |
| **RF** | 0.79 | 0.48 | 0.54 | 0.71 | 0.22 |
| **Window Length – 15** |  |  |  |  |  |
| **ANN** | 0.65 | 0.57 | 0.58 | 0.66 | 0.17 |
| **XGB** | 0.73 | 0.53 | 0.57 | 0.71 | 0.21 |
| **RF** | 0.78 | 0.51 | 0.56 | **0.72** | 0.23 |
| **Window Length – 17** |  |  |  |  |  |
| **ANN** | 0.65 | 0.58 | 0.59 | 0.67 | 0.18 |
| **XGB** | 0.71 | 0.56 | 0.59 | 0.70 | 0.21 |
| **RF** | 0.76 | 0.52 | 0.57 | 0.71 | 0.22 |
| **Window Length – 19** |  |  |  |  |  |
| **ANN** | 0.65 | 0.57 | 0.58 | 0.66 | 0.17 |
| **XGB** | 0.71 | 0.56 | 0.59 | 0.70 | 0.21 |
| **RF** | 0.76 | 0.53 | 0.57 | 0.71 | 0.23 |
| **Window Length – 21** |  |  |  |  |  |
| **ANN** | 0.59 | 0.66 | 0.65 | 0.67 | 0.20 |
| **XGB** | 0.72 | 0.56 | 0.59 | 0.71 | 0.22 |
| **RF** | 0.75 | 0.53 | 0.57 | 0.71 | 0.22 |

### 4.4. Benchmarking

In our study, we observed that Random Forests, trained on Residual Solvent Accessibility (RSA) and Position-Specific Scoring Matrix (PSSM) achieved the highest performance in predicting antibody interacting regions in the antigen. The performance of the ensemble model compared to its individual features is displayed in Figure 6. It is important for any study to benchmark it with existing methods. Best of our knowledge, no method have been developed so far for predicting Conformational B-cell epitopes for human host or for predicting human antibody-interacting residues. Thus direct comparison of our method with previous studies is not possible. In the past, number of methods have been developed for predicting conformational B-cell epitopes not specific to any host. Thus we evaluate these general methods of B-cell epitope prediction on independent dataset used in this study to benchmark these methods with proposed method. The B-cell epitope prediction methods used for benchmarking includes CBTOPE [13], Bepipred-2.0 [37], Bepipred-3.0 [10], SEMA-1d [11], and Epidope [38][. As shown in Table 4, none of the existing methods achieved AUC more than 0.62 on independent dataset, whereas our methods achieved AUC 0.72 on same dataset. Additionally, we also tested HAIRpred on the mouse dataset derived from experimental results through SAbDaB. It is observed that the performance of our methods decrease significantly on mouse dataset. These observation further support to develop host-specific methods for predicting conformational B-cell epitopes.

**Figure 6:**
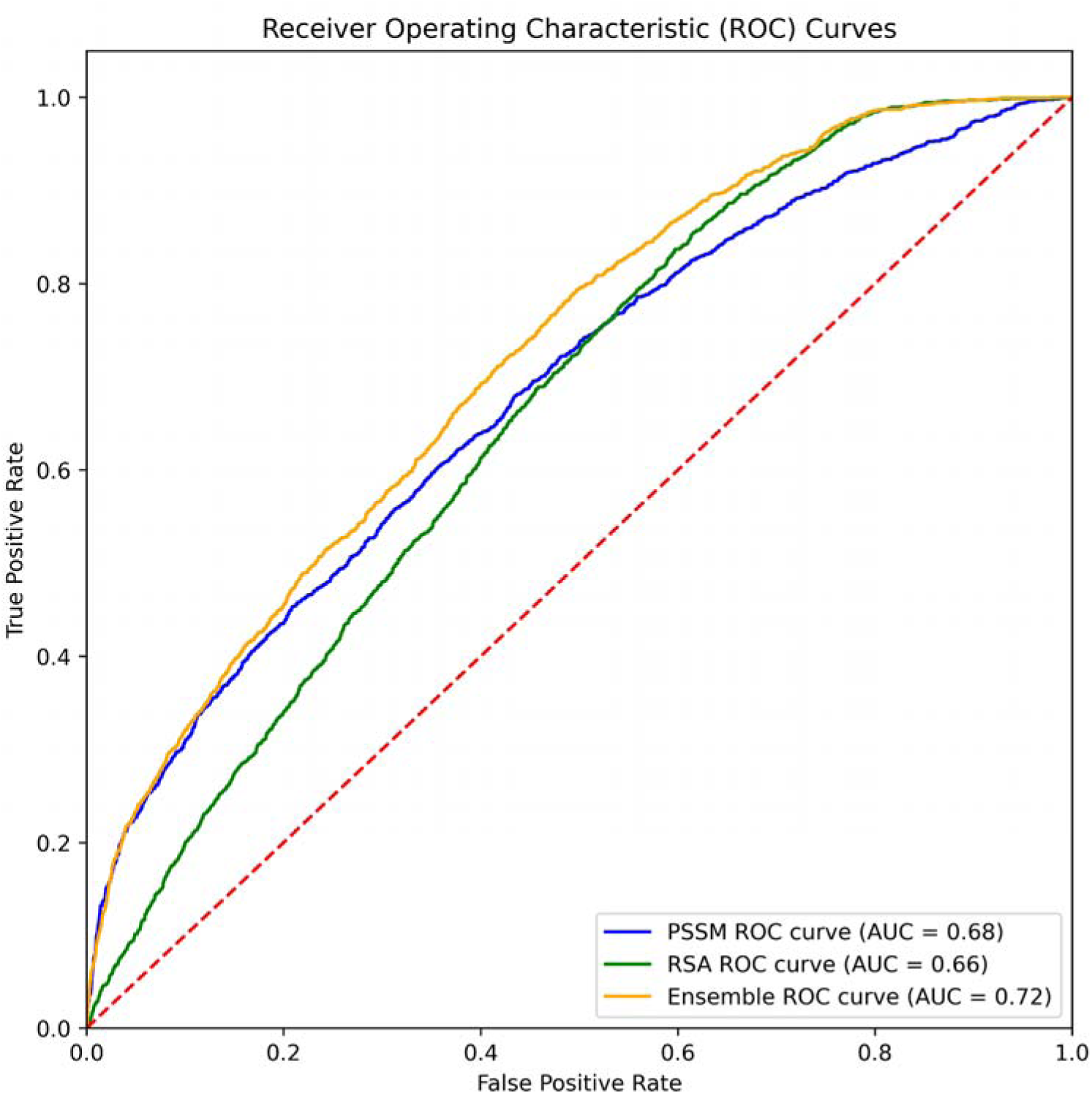
ROC curves of ensemble model (window length = 15) compared with individual features

**Table 4:** The performance of our method HAIRpred and existing methods on independent dataset.

| Model | Year | Sensitivity | Specificity | Accuracy | AUCROC | MCC |
| --- | --- | --- | --- | --- | --- | --- |
| CBTOPE | 2010 | 0.09 | 0.94 | 0.80 | 0.52 | 0.04 |
| Bepipred - 2.0 | 2017 | 0.77 | 0.22 | 0.31 | 0.50 | -0.01 |
| Epidope | 2021 | 0.03 | 0.96 | 0.81 | 0.50 | -0.02 |
| Bepipred - 3.0 | 2022 | 0.65 | 0.58 | 0.59 | 0.62 | 0.17 |
| Sema - 1d | 2024 | 0.65 | 0.60 | 0.61 | 0.62 | 0.18 |
| HAIRpred on Mouse Dataset | -- | 0.71 | 0.54 | 0.56 | 0.68 | 0.17 |
| HAIRpred | -- | 0.78 | 0.51 | 0.56 | 0.72 | 0.23 |

### 4.5. Feature importance analysis

HAIRpred is an ensemble method that combine two Random Forest based models, trained on Residual Solvent Accessibility (RSA) and Position-Specific Scoring Matrix (PSSM) on pattern length 15 respectively. To understand the decision process of these two individual models, Shapley Values were calculated for each model using the python shap package. Figure 7 shows the feature importance by position on the pattern for the RSA trained model. It is observed that the RSA of the central residue in the pattern has the largest influence on the decision making process of the RSA-trained Random Forest Model. Similarly, Shapley Values were calculated for the PSSM-trained model and the top features with highest Shapley Values are : 8Q, 8E, 8R, 8K, 8H, 8N, 8C, 8D, 8A and 11A. Here the number refers to the position of the residue in the pattern and the represents the column of the PSSM matrix. It is similarly observed that the central residue in the pattern has the largest influence on the decision making process for the PSSM-trained model.

**Figure 7:**
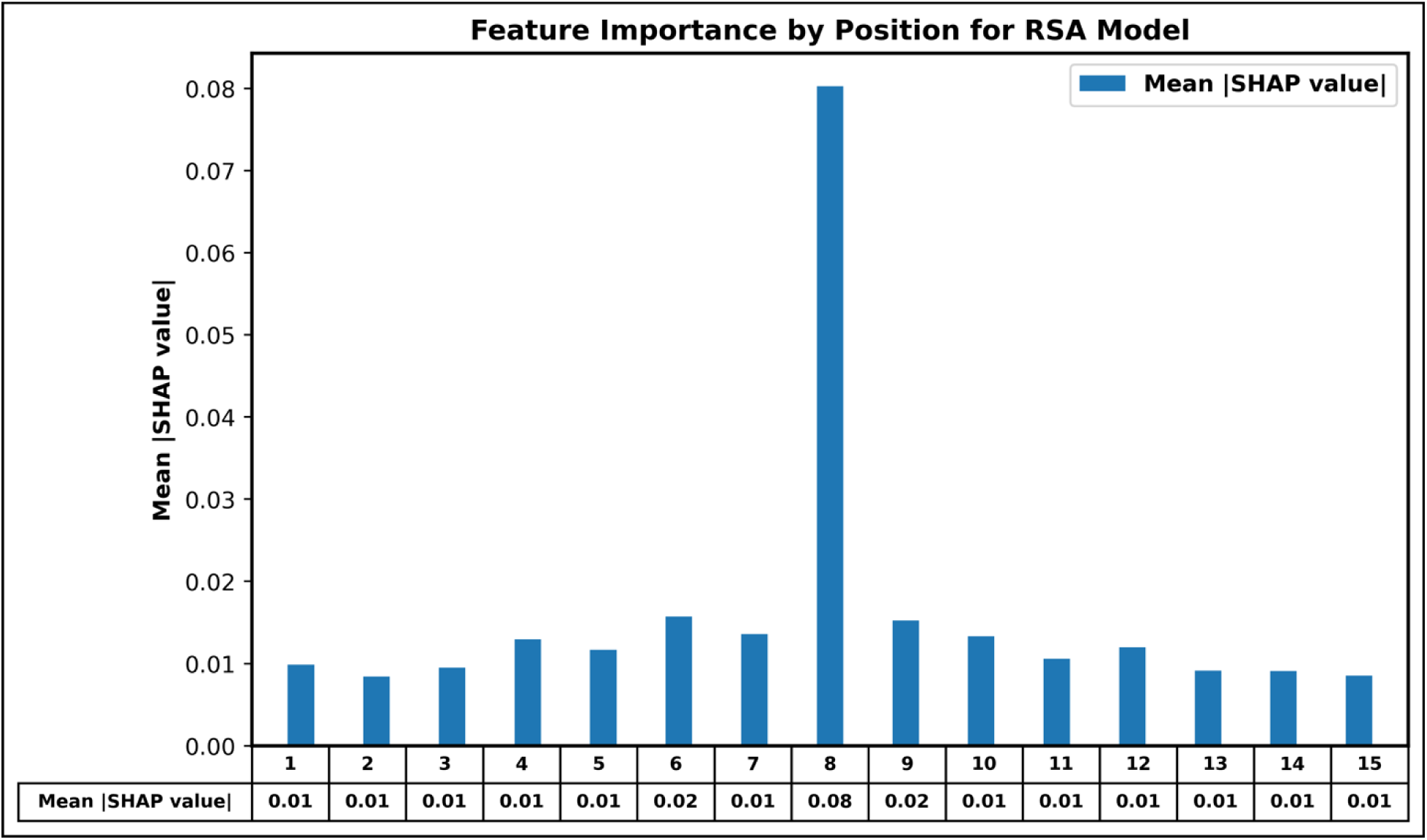
Shapley Values calculated for each position of the pattern for the RSA model

### 4.6. Web Server Implementation

To serve the scientific community, we have developed a user-friendly web server that integrates the best-performing prediction models from our study. This web server provides an accessible platform for researchers to predict antibody-interacting residues in antigen sequences. Users can submit antigen sequences in FASTA format, and the server uses the selected model to predict antibody-binding sites in each submitted antigen. The output is designed to be both extensive and user-friendly, offering results in textual format alongside an intuitive graphical representation of the antibody-interacting residues within the protein or antigen (see Figure 8). In addition to providing predictions, the server allows users to download the training and testing datasets used in this study. The server has been implemented using a combination of HTML, JavaScript, and PHP scripts. The webserver has robust functionality and compatibility across a wide range of devices, including laptops, Android smartphones, iPhones, and iPads. We have also developed a standalone version of HAIRpred to enhance its accessibility. This package offers researchers the flexibility to run predictions locally without the need for active internet connection. The standalone version is available on GitHub (https://github.com/raghavagps/HAIRpred) and can also be installed via PyPI package which is accessible through the pip package (pip install hairpred). Both the web server and standalone package aim to democratize access to advanced computational tools, fostering advancements in antigen-antibody interaction research. The open-source web server can be accessed at https://webs.iiitd.edu.in/raghava/hairpred/, providing a seamless and reliable resource for the scientific community.

**Figure 8:**
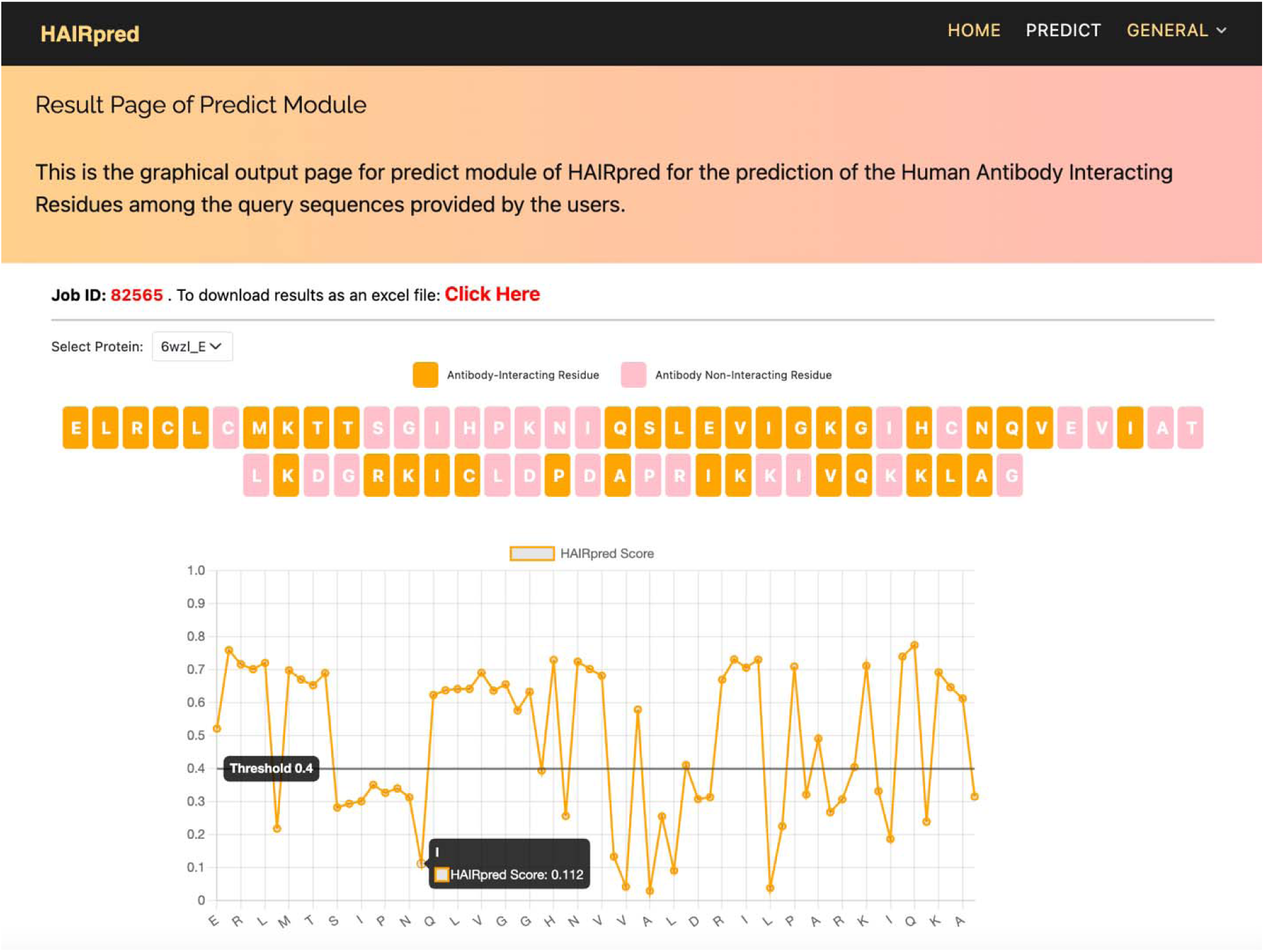
HAIRpred web server interface showing Antibody-Interacting residue prediction results

## 5. Discussion and Conclusion

Antibodies are vital glycoproteins in the adaptive immune system, designed to bind to foreign particles through specific regions called epitopes. Identifying these antibody-binding regions is crucial for advancing immunological research and enabling the development of vaccines, therapeutics, and diagnostic tools. However, existing predictive models face several challenges, including limited host specificity and an inability to account for species-specific variations in immune responses and antibody structures. These shortcomings often lead to suboptimal performance when applied to experimental datasets. Furthermore, many models lack interpretability, offering little understanding of the factors influencing antibody-antigen interactions. To address these limitations, our study focused exclusively on human antibody-antigen complexes derived from experimental data. By narrowing the scope to human systems, we sought to overcome the drawbacks of generalized models that neglect species-specific differences. We curated a comprehensive feature set that includes structural, sequential, and evolutionary characteristics of antigens. Through rigorous testing using data from the Protein Data Bank and the SAbDab database, we identified key features such as Relative Solvent Accessibility (RSA) and Position-Specific Scoring Matrices (PSSM) as the most important determinants of antibody-antigen interactions.

Additionally, we investigated the influence of local antigenic environments by analyzing sequence patterns of varying lengths. This analysis resulted in the development of Human Antibody Interacting Residue Predictor (HAIRpred), an ensemble-based random forest model designed to predict antibody-interacting regions in antigens with high precision. HAIRpred represents a second-generation, host-specific tool that prioritizes human antibody interactions over generalized, cross-species predictions. The model demonstrated a remarkable AUROC of 0.72 on an independent experimental dataset, outperforming existing state-of-the-art models. It also demonstrated superior performance in identifying human antibody-interacting residues compared to mouse-derived antibodies, underscoring the importance of host specificity. Insights from SHAP (SHapley Additive exPlanations) analysis highlighted that central residues in antigenic patterns are pivotal in predicting interactions, enhancing the model’s interpretability. While these results mark significant progress, this study acknowledges one limitation: the sequence similarity threshold of 70% used for data clustering. This threshold was necessary to ensure an adequate amount of training data. However, as more experimental data becomes available, lowering this threshold to 40% will allow a more in-depth exploration of antigen-antibody interactions across diverse antigen structures, further improving the model’s generalizability and performance. In conclusion, HAIRpred offers a major advancement in antibody-antigen interaction prediction by adopting a host-specific, human-centric approach. Its robust performance on human data makes it an invaluable resource for immunological research, with potential applications in vaccine design, therapeutic development, and diagnostics. By addressing the limitations of earlier models, HAIRpred sets a new standard for second-generation, host-specific prediction tools. To ensure accessibility, we have made both a user-friendly web server, PyPI package, and a standalone version of HAIRpred available at https://webs.iiitd.edu.in/raghava/hairpred/.

## Authors’ contributions

RS collected the dataset. RS processed the dataset. RS, NK, and GPSR implemented the algorithms and developed the prediction models. RS, NK, and GPSR analysed the results. RS and NK created the front-end, back-end of the webserver and standalone of the method. RS, NK, and GPSR penned the manuscript. GPSR conceived and coordinated the project. All authors have read and approved the final manuscript.

## Acknowledgments

Authors are thankful to the University Grants Commission (UGC) and DST-Inspire (KVPY), for fellowships and financial support, and the Department of Computational Biology, IIITD New Delhi for infrastructure and facilities. We would like to acknowledge that Figures were created using BioRender.com.

## Data Availability Statement

All the datasets used in this study are available at the “HAIRpred” web server, at https://webs.iiitd.edu.in/raghava/hairpred/.

